# Divorce rate in birds increases with male promiscuity and migration distance

**DOI:** 10.1101/2022.10.13.512018

**Authors:** Yiqing Chen, Xi Lin, Zitan Song, Yang Liu

## Abstract

Socially monogamous animals may break up their partnership after one breeding season by a so-called ‘divorce’ behaviour. Divorce rate immensely varies across avian taxa that have a predominantly monogamous social mating system. Although a range of factors associated with divorce have been tested, there is not a consensus regarding the large-scale variation and relationships among associated factors. Moreover, the impact of sexual roles in divorce still needs further investigation. Here, we applied phylogenetic comparative methods to analyze one of the largest datasets ever compiled that included divorce rates from published case studies of 232 avian species from 25 orders and 61 families. We tested correlations between divorce rate and a group of factors that are closely related to pair bond strength: promiscuity of both sexes, migration distance, and adult mortality. Our results showed that only male promiscuity, but not female promiscuity, had a critical relationship with divorce rate. Furthermore, migration distance was positively correlated with divorce rate and indirectly affected divorce rate via male promiscuity. These findings indicated that divorce might not be simply explained as an adaptive strategy or neutral occurrence, but could be a mixed response to sexual conflict and stress from the ambient environment.

## 1. Introduction

Most avian species form socially monogamous pair bonds, but they may end the bonds because of the death of one partner or ‘remarry’ a different partner after so-called ‘divorce’. Divorce can be defined as an individual re-mating with a new partner while its former partner of the last breeding season is still alive.^1,2^ Divorce involves breaking up a pair bond and re-selecting a new mate; this is linked to sexual selection and plays an important role in individual fitness^2,3,23,24^. Compared with long-term partnership, individuals mate with different partners and create novel genetic variation in offspring^4,22,25,26^; thus, divorce may be a mating strategy that impacts population dynamics and promotes intra-specific gene flow^22,25–27^. There are two main hypotheses on causes of divorce. One explains divorce as an adaptive strategy that boosts individual reproductive fitness, whereas the other indicates that divorce is neutral or an indirect effect of other ecological drivers, such as mortality and migration (Table 1). ^1,5,6^

**Table 1.**
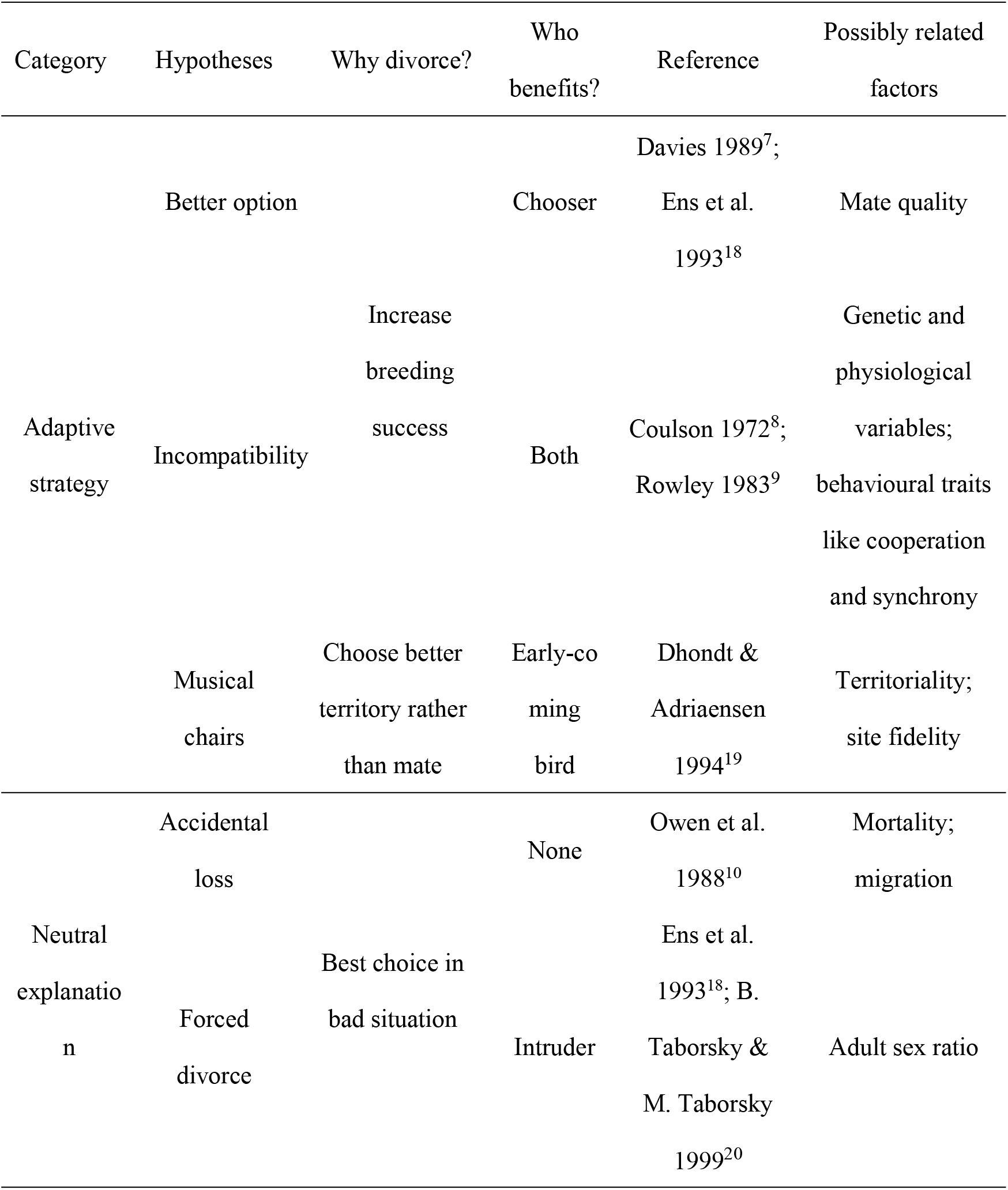
Summary of the major hypotheses regarding divorce in birds.

A range of factors associated with divorce rate have been documented in case studies, including mortality^2,11^, migration^2,12^, adult sex ratio^13^, and extra-pair paternity^14,21,22^. However, these studies only cover a limited range of avian species. Consensus regarding explanations for the global-scale variation of divorce rate is still lacking, and relationships among multiple factors of divorce remain unclear. Among hypotheses, predicted benefits vary for either member of a pair, and it remains unclear which sex benefits from divorce. Because there are obvious differences in fitness consequences for males and females in a single reproductive event, sex-specific roles in divorce should be expected.

To better address these issues, we compiled a large dataset of divorce rate for 232 avian species from 25 orders and 61 families, and some correlates such as sex-specific promiscuity, migration distance, and adult annual mortality (for details, see Methods), which are factors closely related to pair bond strength. Our dataset includes both geography and phylogeny (Figure 1). Here, we used phylogenetic comparative methods to test the following hypotheses: (1) high promiscuity in either sex predicts high divorce rate; (2) longer migration distance may increase divorce rate through asynchrony; and (3) higher mortality rate lowers the likelihood that a partner will reunite with its partner, and thus increases divorce rate. In addition, we conducted phylogenetic path analyses (PPA) to elucidate the unknown relationships among correlates to better understand potential indirect effects on divorce rate.

**Figure 1.**
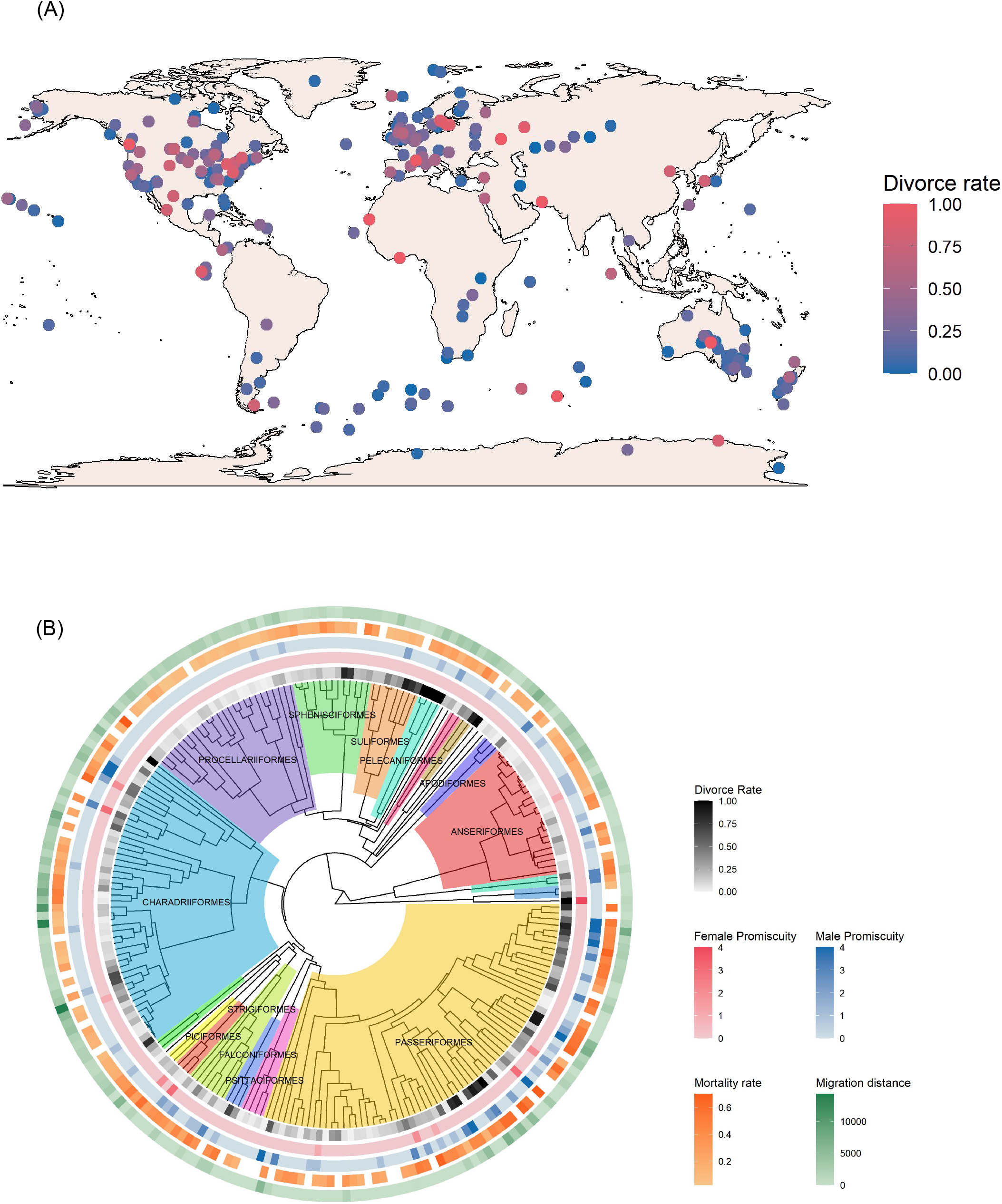
Geographic and phylogenetic distributions of divorce rate in birds. (A) Geographic distribution of divorce rate in birds (n = 232 species). Species presence was mapped as study sites of each case study or as centroid coordinates from AVONET (Tobias et al., 2022) if the data source did not provide specific locations. (B) Phylogenetic distribution of divorce rate, promiscuity score, mortality rate, and migration distance in birds (n = 232 species). Male and female promiscuity were scored as (0) no (or very rare) polygamy (< 0.1%), (1) rare polygamy (0.1%–1%), (2) uncommon polygamy (1%–5%), (3) moderate polygamy (5%–20%), and (4) common polygamy (> 20%) (for details, see Methods). Migration distance is shown in metres. Clade colours represent different taxonomic orders.

## 2. Methods

### (a) Divorce rate

We used data from Kenny et al. (2017)^15^, Liker et al. (2014)^13^, Botero et al. (2012)^16^, *Handbook of the Birds of the World* (https://birdsoftheworld.org)^17^, and other published literature^18–56^. Annual divorce rate was defined as the percentage of pairs that both survived but changed mates from one year to the next year in a population and was only measured in monogamous pairs. For multiple reports in one species, we calculated the average of the reported data.

### (b) Female and male promiscuity measures

Promiscuity scores were used to reflect the mating system variation for both sexes. Our study only involved species that predominantly exhibit monogamy because divorce only applies to socially monogamous species. Some of these socially monogamous species still have a proportion of polygamy or polygynandry described in *Handbook of the Birds of the World* (https://birdsoftheworld.org)^17^. We considered the proportion of polygamy as a measure of the potential for either sex to have more mates. These promiscuity scores were based on the description from *Handbook of the Birds of the World* as follows:

1. 0 for < 0.1% polygamous/polygynandrous individuals, or just the key word “monogamous” appearing with no detailed description indicated that there are rare exceptions of polygamy or polygynandry.
2. 1 for 0.1%–1% polygamous/polygynandrous individuals, or those with the key words “(permanently/predominately/primarily/usually/…) monogamous” and “(extremely rare/occasional/…) polygamy/polygynandry”, which indicates that polygamy/polygynandry are not the primary mating system, or some detailed description indicating that there are rare exceptions of polygamy/polygynandry.
3. 2 for 1%–5% polygamous/polygynandrous individuals, or those with the key words “(permanently/predominately/primarily/usually/…) monogamous” and the occurrence of polygamy/polygynandry was higher than that for score 1 but closer to score 1 than score 4.
4. 3 for 5%–20% polygamous/polygynandrous individuals, or those with the key word “polygamy”/“polygynandry”, even when “monogamous” appears, but the occurrence of polygamy/polygynandry was lower than score 4 and labile.
5. 4 for > 20% polygamous/polygynandrous individuals, or those with the key word “polygamy”/“polygynandry” even when “monogamous” appears or a detailed description such as “males/females mate with multiple partners”.

### (c) Other traits

For migration distance, we used data from Delhey et al. (2021)^57^. Adult mortality rate was extracted from the AVONET database^58^. Our final dataset contained 232 avian species from 25 orders and 61 families, of which the 186 species had a complete dataset.

### (d) Phylogenetic analyses

To control phylogenetic uncertainty, we used 100 randomly selected phylogenetic trees extracted from birdtree.org^59^. We ran a full model containing all four predictors (male and female promiscuity, mortality rate, and migration distance) of divorce rate on a subset of 186 species using the MCMCglmm procedure in R^60^ version 4.2.1. We used the priors [list(R=list(V=1, nu=0.002), G=list(G1=list(V=1, nu=1, alpha.mu=0, alpha.V=1000)))] and ran the MCMC algorithm for 75,000 iterations, with thinning of 40 and burn-in of 7,500.

The model was based on 186 avian species and was generated and implemented in the R package ‘MCMCglmm’^61^. The phylogenetic effects were based on 100 Hackett backbone trees from birdtree.org. Migration distance, and male and female promiscuity scores were log_10_-transformed and scaled. Significant counts referred to the presence of significant p values in 100 iterations.

To determine the robustness of our results, we also tested the same model using the Phylogenetic Generalized Least Squares (PGLS) approach in the R package ‘caper’^62^ and estimated the phylogenetic signal by optimizing the λ parameter. In this procedure, we assumed that promiscuity scales reflected continuous variation in the degree of polygamy.

To inspect possible direct and indirect relationships among the five traits, we conducted PPA, which uses phylogenetic independent contrasts and allows testing of alternative models by determining the path coefficients and overall model fit. Our 95 candidate models contained all potential combinations of hypothesized relationships among traits. We ran the analyses and estimated the best-supported model using the R package ‘phylopath’^63^. Standardized regression coefficients of the path were considered statistically significant when 95% confidence intervals did not include zero.

Model codes correspond to diagrams presented in Figure S1. For each model, we reported the C-statistic (C), p-value, CICc value, ΔCICc value, and CICc weights (ω). P-values of the C-statistic were used to determine significance and indicate if the model was rejected by the data. Models were based on 186 avian species. Results from all tested models are provided in Table S2.

All data quantification, analysis, and visualization were conducted in RStudio^64^ version 2022.07.0+548 and R^60^ version 4.2.1.

## 3. Results

MCMCglmm results for 100 random trees all showed that male promiscuity had significant and positive correlations with divorce rate (Table 2; MCMCglmm, estimate [SE] = 0.0570 [0.0161], p < 0.001, n = 186 species), which indicated that species with higher proportions of male polygamy have higher divorce rates in monogamous pairs. In contrast, female promiscuity did not show any significant effect on divorce rate (Table 2; MCMCglmm, Estimate [SE] = −0.0080 [0.0160], p > 0.05, n = 186 species) in all iterations of 100 random trees. PGLS analyses had similar results (Table S1), which to some extent supported the robustness of our results.

**Table 2.** Effects of male and female promiscuity, migration distance, and mortality rate on divorce rate across 186 bird species.

|  | Estimate | Estimate 95% CI | SE | z value | p-value | Significant count |
| --- | --- | --- | --- | --- | --- | --- |
| Intercept | 0.1963 | -0.041 — 0.431 | 0.1206 | 1.6281 | 0.1033 | 9 |
| Male promiscuity | 0.0566 | 0.025 — 0.088 | 0.0161 | 3.5167 | **0.0005** | 100 |
| Female promiscuity | -0.0080 | -0.039 — 0.023 | 0.0160 | -0.4977 | 0.6314 | 0 |
| Migration distance | 0.0476 | 0.011 — 0.084 | 0.0186 | 2.5610 | **0.0104** | 100 |
| Mortality rate | 0.1626 | -0.082 — 0.404 | 0.1241 | 1.3110 | 0.1911 | 0 |

Results in 100 random trees all showed a significant and positive correlation between migration distance and divorce rate (Table 2; MCMCglmm, estimate = 0.0476 [0.0186], p < 0.05, n = 186 species), which indicates that species with longer migration distances had higher divorce rates. However, mortality rate did not show any significant effect on divorce rate (Table 2; MCMCglmm, p > 0.05, n = 186 species) in any iterations of 100 random trees, which seems to contradict previous opinions^5^. PGLS analyses also showed similar results (Table S1), which further supported our findings.

The best-supported PPA model (mean CICc = 33.6987) with average standardized regression coefficients(Figure 2, Table S2) indicated that there was no direct effect of female promiscuity on divorce rate, which was consistent with our MCMCglmm results. Female promiscuity was only affected by male promiscuity, whereas mortality rate was only affected by migration distance in the path model. Although mortality rate showed no direct relationship with divorce, it might indirectly raise divorce rate via male promiscuity. Moreover, migration distance can both directly affect divorce rate and indirectly affect divorce rate through male promiscuity. Longer migrants had an increased trend of mortality, male promiscuity, and divorce rate.

**Figure 2.**
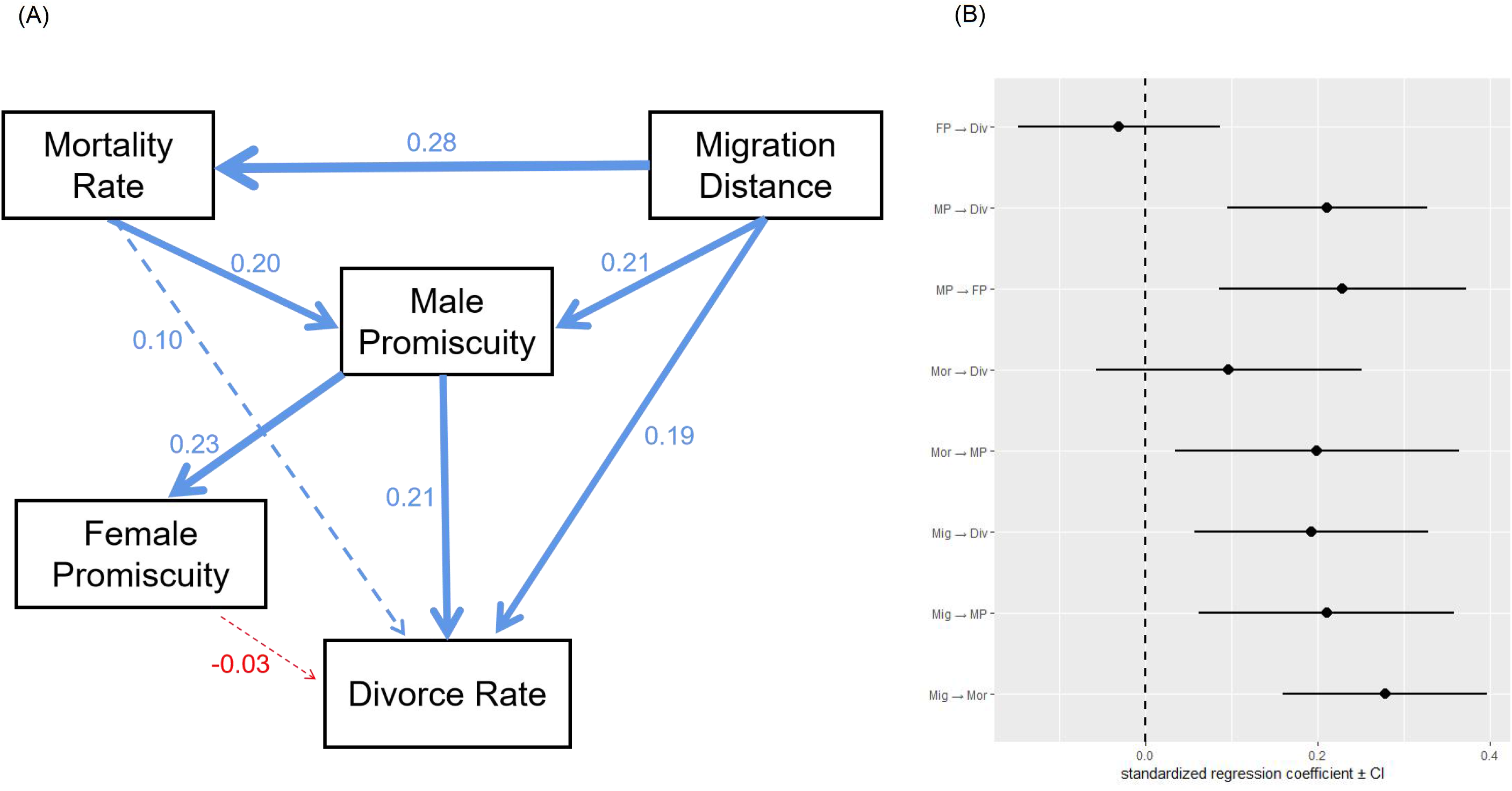
Phylogenetic path model showing how promiscuity of both sexes, mortality, and migration distance affect divorce rate in birds (n = 186 species). (A) Path diagram showing the best-supported models by the data (Table S2). Arrows indicate direct effects; arrow colour indicates the direction of the effect (blue, positive; red, negative). The absolute value of the standardized regression coefficient (Figure S1) is indicated by numeric values and line widths (higher values have wider lines). Solid lines indicate significant relationships and dotted lines indicate non-significant relationships. (B) Standardized regression coefficients for the path models. The predictor variables were scaled. The centre point denotes the mean and the bars denote the 95% lower and upper confidence limits calculated by model-averaging. A standardized regression coefficient was considered statistically significant when 95% confidence intervals did not include 0. Abbreviations: Div, divorce rate; MP, male promiscuity score; FP, female promiscuity score; Mor, mortality rate; Mig, migration distance.

## 4. Discussion

Through a combined approach of phylogenetic comparative methods and path analyses, this study had three key findings. First, male promiscuity rather than female promiscuity raised divorce rate. Second, migration distance was positively associated with divorce rate and might also affect divorce rate through male promiscuity. Finally, we found no evidence for a direct relationship between mortality and divorce rate.

Although divorce rate is only defined in socially monogamous birds, some socially monogamous species still show a certain proportion of polygamy or polygynandry according to *Handbook of the Birds of the World* (male polygamy is mentioned for 62 species and female polygamy for 12 species; the rest of the species had no description of polygamy), and these descriptions were clearly distinct from those of extra-pair paternity. In this study, we measured the amount of “promiscuity” based on these descriptions, which could be considered a measurement of the variation of mating systems in certain populations. Cézilly and Nager (1995)^14^ discovered a positive correlation between divorce rate and extra-pair paternity, but it was difficult to separate the effects of the two sexes using extra-pair paternity data, because it only showed the paternity of the offspring while which sex initiated extra-pair copulation remained unknown. In this study, we considered variation of promiscuity behaviour in the different sexes and revealed different effects of the two sexes on divorce rate.

Our results showed that only the proportion of male promiscuity in the population raised divorce rate. It is thought that there is relatively less investment in breeding for males than females^65^; therefore, males may be less adversely impacted by divorce. Variation in mating system may improve fitness of males as they reduce their fidelity to a single female^65^, and divorce further contributes to this strategy. Liker et. al (2014)^13^ reported that divorce is more frequent in species or populations with a female-biased sex ratio because males have more available potential mates, which may be consistent with our results. However, females have larger costs in breeding, so they must be more prudent in choosing partners. Unlike males that can afford to correct errors, females rely on former breeding experience as a more robust strategy. Therefore, females tend to maintain an old pair bond rather than divorce, even though they have some chance to be polygamous. However, some studies indicated that divorce only benefits female fitness^23^ and that females initiate divorce in certain species^28^, which contradict our results.

As expected, our results confirmed that divorce rates were higher in species with longer migration distances. Studies have revealed that divorce rates are higher in migratory species than resident species^2,12^. However, we used specific migration distance in analyses rather than simple classification such as resident, semi-resident, and migrant. Thus, our results expand the conclusion that both migration occurrence and distance affect divorce rate. This result could be explained by asynchrony in migration^29–34^, as longer migration might amplify the time-lag of arrival between partners and lead to a higher degree of asynchrony in arrival. Moreover, long-distance migration extends travel time and narrows the time window for breeding. In this context, divorce could be a salvage strategy to ensure breeding for the year when partner do not arrive with each other, which was previously shown in some long-distance migratory waterbirds^34–36^.

In addition, PPA results showed a positive effect of migration distance on male promiscuity, which is consistent with a previous study that showed long-distance migrant species have larger testes and a greater tendency to seek more mating chances than resident species^59^. Thus, it is possible that migration distance may also indirectly raise divorce rate through male promiscuity. However, the relationship between migration distance and divorce rate might not just be a simple positive correlation. When migration distance was scored as residents, short-distance migrants, variable migrants, and long-distance migrants (Figure S2), the variable migrants had the highest divorce rate, rather than the long-distance migrants. This pattern is possibly related to mechanisms of how synchrony is achieved in migratory birds^29^. It is possible that for short-distance migrants, asynchrony can be mainly explained by incompatibility, and divorce could be an adaptive strategy; alternatively, for long-distance migrants, asynchrony is mostly affected by environmental conditions and divorces are forced. Variable migrants may have mixed influences and both effects collectively increase divorce rate.

Our PPA results indicated that divorce could be influenced by both subjective (male promiscuity) and objective factors (migration). Divorce might not be a simple adaptive or non-adaptive strategy, but could be simultaneously affected by the decisions of the sexes and stress from the environment. However, we only found indirect rather than direct correlations between mortality and divorce rate. It is possible that mortality and divorce rate showed a co-varying pattern that was influenced by both migration distance and male promiscuity.

However, there are also limitations of our work. PGLS results showed relatively high phylogenetic signal (λ=0.78, Table S1), which indicated that divorce behaviour was driven by phylogenetic constraints to a certain extent. Moreover, our study did not consider trait variation within populations. Finally, our dataset, especially for divorce and mortality, was too limited to represent the entire avian tree of life. A larger dataset that includes continuous studies and more advanced theoretical modeling research might help us gain an even better understanding of the general drivers of bird divorce.

## Ethic statement

This research only used data from published literature, without involving human subjects. We did not use any animals, DNA, or fossil samples.

## Data accessibility

Data are available from the Dryad Digital Repository: Chen, Yiqing; Lin, Xi; Song, Zitan; Liu, Yang (2022), Avian Divorce rate, Dryad, Dataset, https://doi.org/10.5061/dryad.cvdncjt75

## Authors’ contributions

Y.Q. C.: formal analysis, visualization, writing—original draft and writing—review and editing; X. L.: data curation, methodology, and writing—review and editing; Z. T. S.: conceptualization, methodology, formal analysis, investigation, validation, and writing—review and editing; Y. L.: conceptualization, funding acquisition, project administration, resources, supervision, validation, and writing—review and editing. All authors gave final approval for publication and agreed to be held accountable for the work performed therein.

## Acknowledgements

We thank Yuqing Han and Dan Liang for thegir advice on result visualization. We thank Mallory Eckstut, PhD, from Liwen Bianji (Edanz) (www.liwenbianji.cn) for editing the English text of a draft of this manuscript.

