## Supplemental Files for "Divorce rate in birds increases with male promiscuity and migration distance"

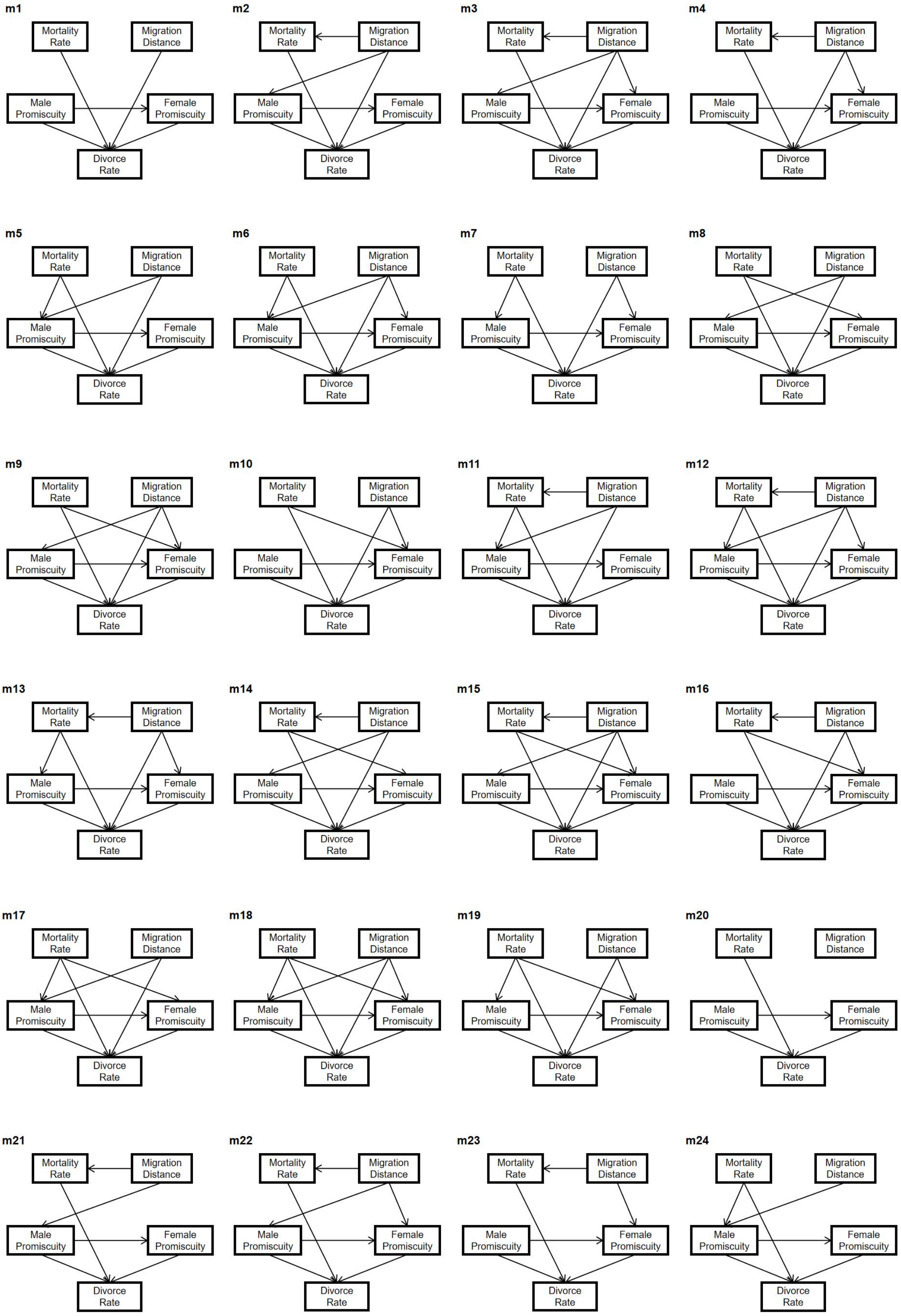

m1
Mortality Rate
Migration Distance
Male Promiscuity
Female Promiscuity
Divorce Rate
m2
Mortality Rate
Migration Distance
Male Promiscuity
Female Promiscuity
Divorce Rate
m3
Mortality Rate
Migration Distance
Male Promiscuity
Female Promiscuity
Divorce Rate
m4
Mortality Rate
Migration Distance
Male Promiscuity
Female Promiscuity
Divorce Rate
m5
Mortality Rate
Migration Distance
Male Promiscuity
Female Promiscuity
Divorce Rate
m6
Mortality Rate
Migration Distance
Male Promiscuity
Female Promiscuity
Divorce Rate
m7
Mortality Rate
Migration Distance
Male Promiscuity
Female Promiscuity
Divorce Rate
m8
Mortality Rate
Migration Distance
Male Promiscuity
Female Promiscuity
Divorce Rate
m9
Mortality Rate
Migration Distance
Male Promiscuity
Female Promiscuity
Divorce Rate
m10
Mortality Rate
Migration Distance
Male Promiscuity
Female Promiscuity
Divorce Rate
m11
Mortality Rate
Migration Distance
Male Promiscuity
Female Promiscuity
Divorce Rate
m12
Mortality Rate
Migration Distance
Male Promiscuity
Female Promiscuity
Divorce Rate
m13
Mortality Rate
Migration Distance
Male Promiscuity
Female Promiscuity
Divorce Rate
m14
Mortality Rate
Migration Distance
Male Promiscuity
Female Promiscuity
Divorce Rate
m15
Mortality Rate
Migration Distance
Male Promiscuity
Female Promiscuity
Divorce Rate
m16
Mortality Rate
Migration Distance
Male Promiscuity
Female Promiscuity
Divorce Rate
m17
Mortality Rate
Migration Distance
Male Promiscuity
Female Promiscuity
Divorce Rate
m18
Mortality Rate
Migration Distance
Male Promiscuity
Female Promiscuity
Divorce Rate
m19
Mortality Rate
Migration Distance
Male Promiscuity
Female Promiscuity
Divorce Rate
m20
Mortality Rate
Migration Distance
Male Promiscuity
Female Promiscuity
Divorce Rate
m21
Mortality Rate
Migration Distance
Male Promiscuity
Female Promiscuity
Divorce Rate
m22
Mortality Rate
Migration Distance
Male Promiscuity
Female Promiscuity
Divorce Rate
m23
Mortality Rate
Migration Distance
Male Promiscuity
Female Promiscuity
Divorce Rate
m24
Mortality Rate
Migration Distance
Male Promiscuity
Female Promiscuity
Divorce Rate

**m25**

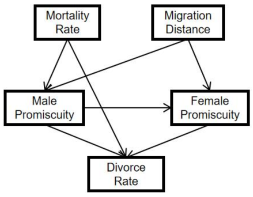


**m26**

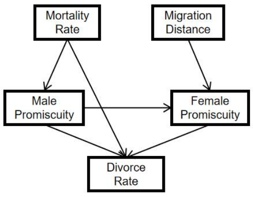


**m27**

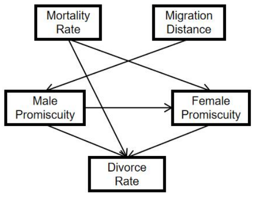


**m28**

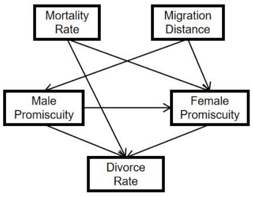


**m29**

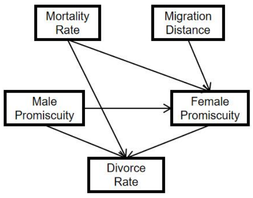


**m30**

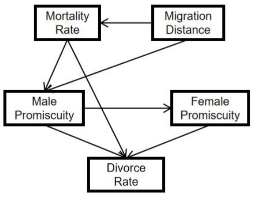


**m31**

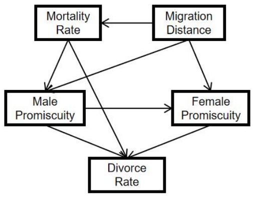


**m32**

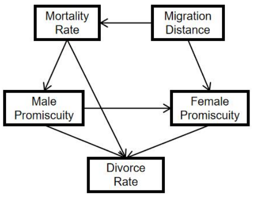


**m33**

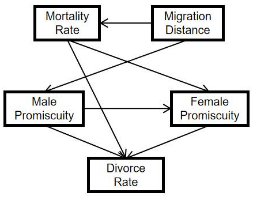


**m34**

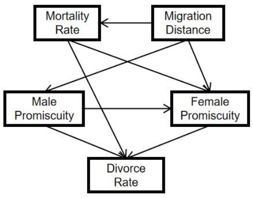


**m35**

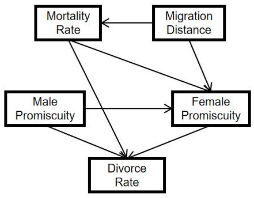


**m36**

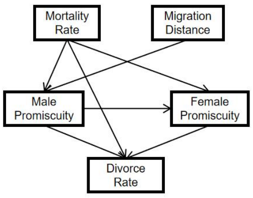


**m37**

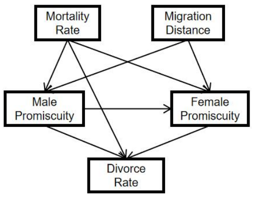


**m38**

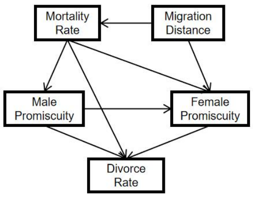


**m39**

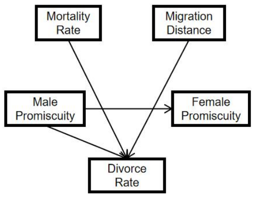


**m40**

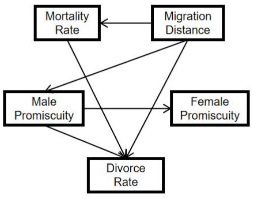




**m41**

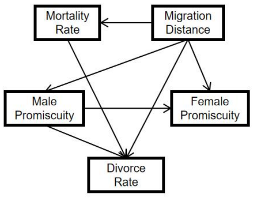


**m42**

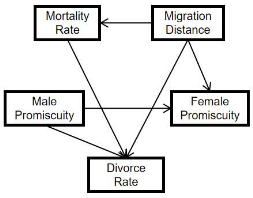


**m43**

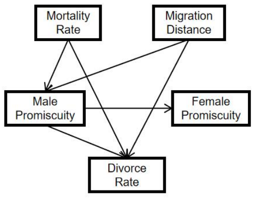


**m44**

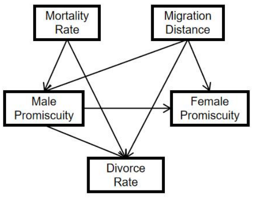


**m45**

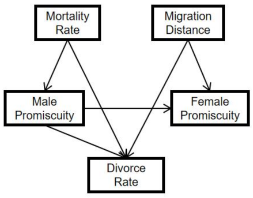


**m46**

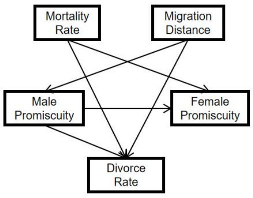


**m47**

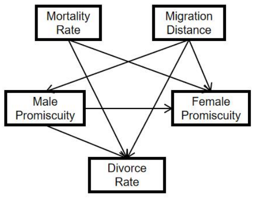


**m48**

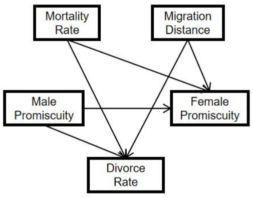

**m49**

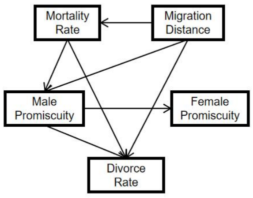


**m50**

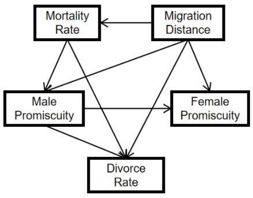


**m51**

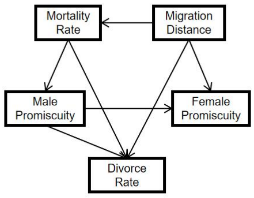


**m52**

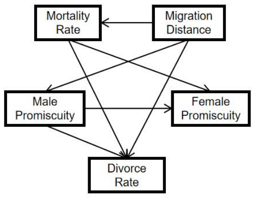


**m53**

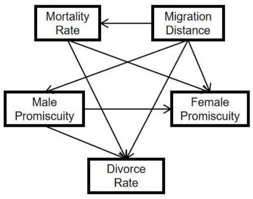


**m54**

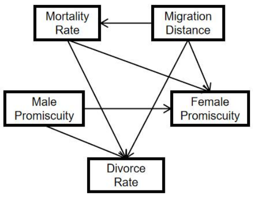


**m55**

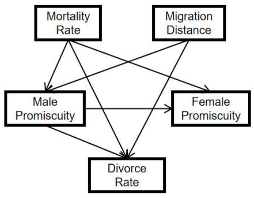


**m56**

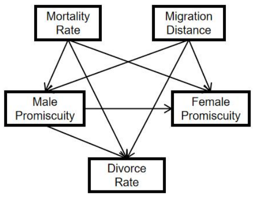


8

**m57**

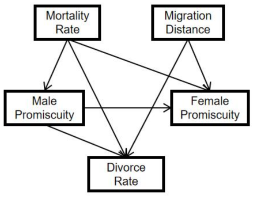


**m58**

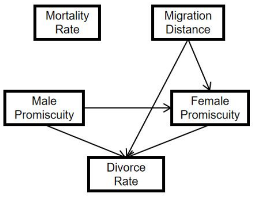


**m59**

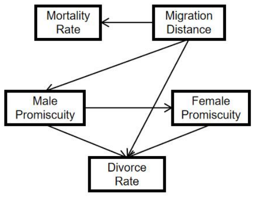


**m60**

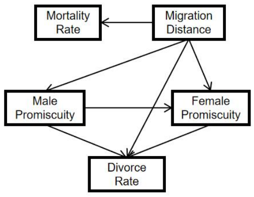


**m61**

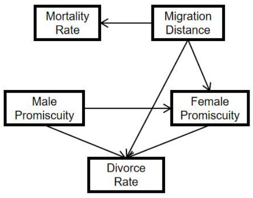


**m62**

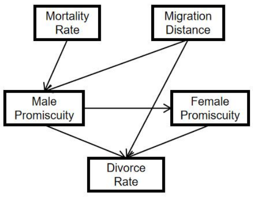


**m63**

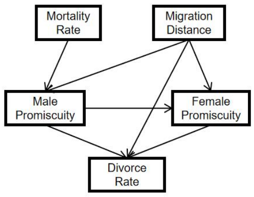


**m64**

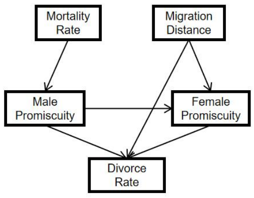


**m65**

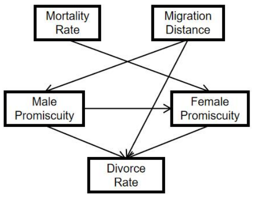


**m66**

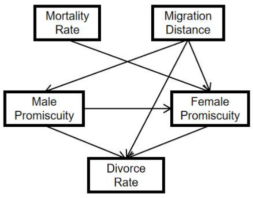


**m67**

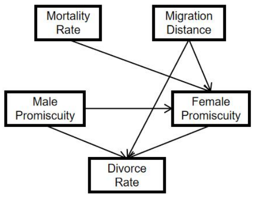


**m68**

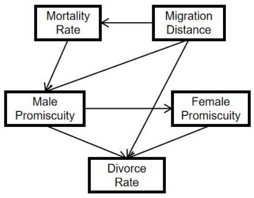


**m69**

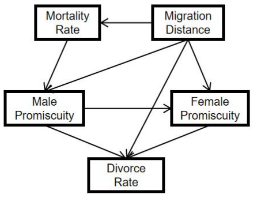


**m70**

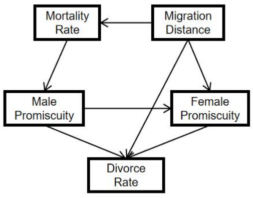


**m71**

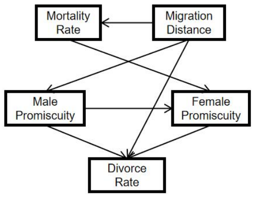


**m72**

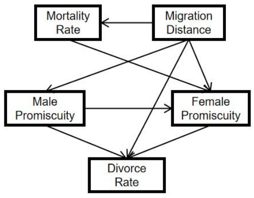


10

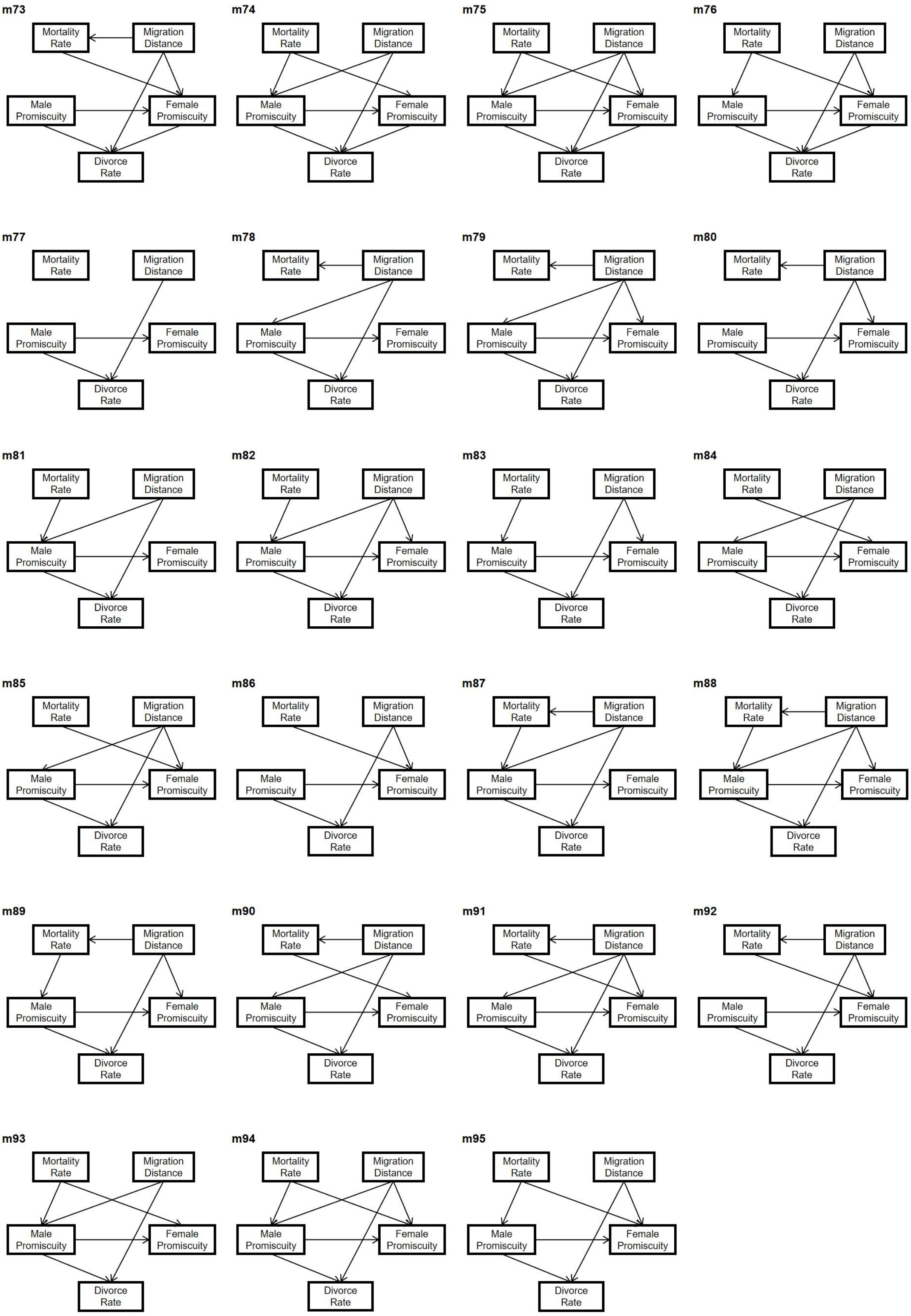


**Figure S1 Path diagrams for all tested path models.** Arrows indicate hypothesized direct relationships between factors.

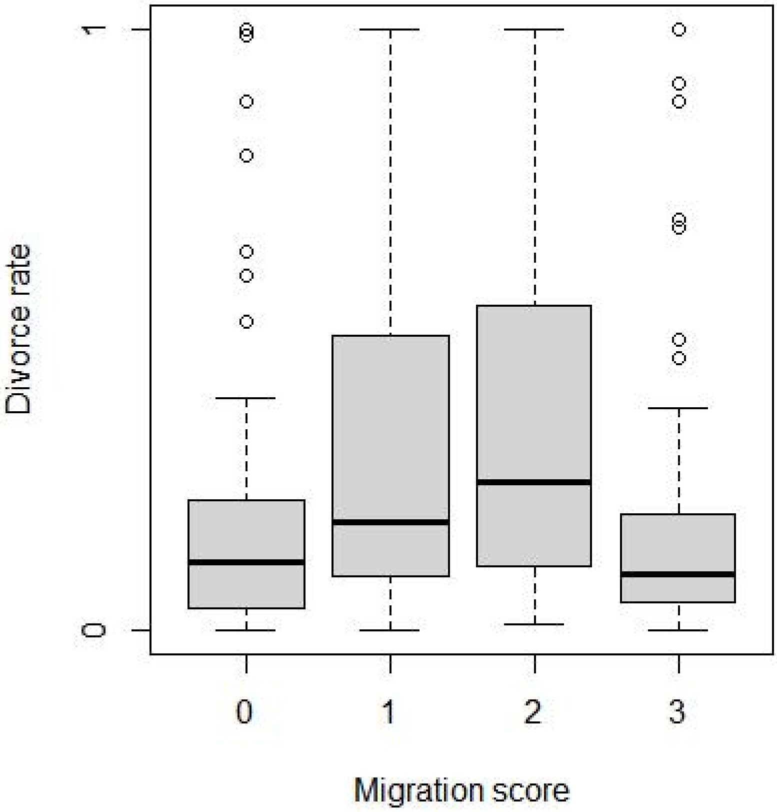


**Figure S2 Migration score and divorce rate.** Migration score was used from Delhey et al., classified as (0) residents, (1) short-distance migrants (migrate < 2,000 km), (2) variable migrants (some populations migrate more than > 2,000 km and some less) and (3) long-distance migrants (> 2,000 km). Central lines represent median values, the top and bottom lines of the box represent the first and third quartiles and circles denote outliers.

**Table S1 The effect of male and female promiscuity, migration distance and mortality rate on divorce rate (evaluated by PGLS)**

| | Estimate 95% CI | p-value 95% CI | $\lambda$ 95% CI |
|---|---|---|---|
| Model | | **0.0000 — 0.0000** | 0.7804 — 0.7885 |
| Male promiscuity | 0.0562 — 0.0569 | **0.0005 — 0.0006** | |
| Female promiscuity | -0.0081 — -0.0078 | 0.6107 — 0.6276 | |

| | | |
|---|---|---|
| Migration distance | 0.0475 — 0.0483 | **0.0103 — 0.0116** |
| Mortality rate | 0.1591 — 0.1631 | 0.1898 — 0.2011 |

Phylogenetically corrected least-squares analyses (PGLS) using 100 iterations. The model was based on 186 avian species and was fitted using package "pgls". The phylogenetic signal for continuous and ordinal variable is measured by Pagel’s lambda (λ), the higher the transition rates, the higher the phylogenetic signal. Migration distance, male and female promiscuity scores were log-10 or square transformed and scaled (detail see Methods). Estimate and p-value of probability of PGLS are shown. Significant p values are highlighted in bold.

**Table S2 Ranking of the best path models from all candidate models based on CICc values**

| model | k | q | C | p-value | CICc | ΔCICc | l | ω |
|---|---|---|---|---|---|---|---|---|
| **m49** | **3** | **12** | **7.3409** | **0.2905** | **33.1443** | **0.0000** | **1.0000** | **0.2054** |
| **m87** | **4** | **11** | **9.9562** | **0.2681** | **33.4734** | **0.3291** | **0.8483** | **0.1743** |
| **m11** | **2** | **13** | **6.3622** | **0.1737** | **34.4785** | **1.3342** | **0.5132** | **0.1054** |
| m50 | 2 | 13 | 7.1824 | 0.1266 | 35.2987 | 2.1544 | 0.3406 | 0.0700 |
| m68 | 3 | 12 | 9.5091 | 0.1469 | 35.3125 | 2.1682 | 0.3382 | 0.0695 |
| m88 | 3 | 12 | 9.7977 | 0.1334 | 35.6012 | 2.4568 | 0.2928 | 0.0601 |
| m52 | 3 | 12 | 10.1865 | 0.1170 | 35.9900 | 2.8457 | 0.2410 | 0.0495 |
| m12 | 1 | 14 | 6.2038 | 0.0450 | 36.6599 | 3.5156 | 0.1724 | 0.0354 |
| m90 | 4 | 11 | 13.1512 | 0.1068 | 36.6685 | 3.5241 | 0.1717 | 0.0353 |
| m53 | 2 | 13 | 8.9154 | 0.0633 | 37.0316 | 3.8873 | 0.1432 | 0.0294 |
| m14 | 2 | 13 | 9.2079 | 0.0561 | 37.3242 | 4.1798 | 0.1237 | 0.0254 |
| m69 | 2 | 13 | 9.3506 | 0.0529 | 37.4669 | 4.3226 | 0.1152 | 0.0237 |
| m91 | 3 | 12 | 11.8801 | 0.0647 | 37.6835 | 4.5392 | 0.1034 | 0.0212 |
| m71 | 3 | 12 | 12.3547 | 0.0545 | 38.1582 | 5.0139 | 0.0815 | 0.0167 |
| m15 | 1 | 14 | 7.9368 | 0.0189 | 38.3929 | 5.2486 | 0.0725 | 0.0149 |
| m40 | 4 | 11 | 15.2776 | 0.0540 | 38.7949 | 5.6505 | 0.0593 | 0.0122 |
| m78 | 5 | 10 | 17.8929 | 0.0568 | 39.1500 | 6.0057 | 0.0496 | 0.0102 |
| m72 | 2 | 13 | 11.0836 | 0.0256 | 39.1999 | 6.0555 | 0.0484 | 0.0099 |
| m2 | 3 | 12 | 14.2990 | 0.0265 | 40.1025 | 6.9581 | 0.0308 | 0.0063 |
| m41 | 3 | 12 | 15.1192 | 0.0193 | 40.9226 | 7.7783 | 0.0205 | 0.0042 |
| m59 | 4 | 11 | 17.4458 | 0.0258 | 40.9631 | 7.8187 | 0.0201 | 0.0041 |
| m79 | 4 | 11 | 17.7345 | 0.0233 | 41.2517 | 8.1074 | 0.0174 | 0.0036 |
| m30 | 3 | 12 | 15.7104 | 0.0154 | 41.5139 | 8.3696 | 0.0152 | 0.0031 |
| m3 | 2 | 13 | 14.1405 | 0.0069 | 42.2568 | 9.1125 | 0.0105 | 0.0022 |
| m60 | 3 | 12 | 17.2874 | 0.0083 | 43.0908 | 9.9465 | 0.0069 | 0.0014 |
| m51 | 3 | 12 | 17.4295 | 0.0078 | 43.2330 | 10.0887 | 0.0064 | 0.0013 |

| | | | | | | | | |
|---|---|---|---|---|---|---|---|---|
| m89 | 4 | 11 | 20.0448 | 0.0102 | 43.5621 | 10.4177 | 0.0055 | 0.0011 |
| m31 | 2 | 13 | 15.5520 | 0.0037 | 43.6683 | 10.5239 | 0.0052 | 0.0011 |
| m33 | 3 | 12 | 18.5561 | 0.0050 | 44.3596 | 11.2152 | 0.0037 | 0.0008 |
| m13 | 2 | 13 | 16.4509 | 0.0025 | 44.5672 | 11.4229 | 0.0033 | 0.0007 |
| m70 | 3 | 12 | 19.5977 | 0.0033 | 45.4012 | 12.2569 | 0.0022 | 0.0004 |
| m34 | 2 | 13 | 17.2849 | 0.0017 | 45.4012 | 12.2569 | 0.0022 | 0.0004 |
| m21 | 4 | 11 | 23.6472 | 0.0026 | 47.1644 | 14.0201 | 0.0009 | 0.0002 |
| m22 | 3 | 12 | 23.4887 | 0.0006 | 49.2922 | 16.1479 | 0.0003 | 0.0001 |
| m54 | 3 | 12 | 23.5390 | 0.0006 | 49.3425 | 16.1981 | 0.0003 | 0.0001 |
| m92 | 4 | 11 | 26.5037 | 0.0009 | 50.0209 | 16.8766 | 0.0002 | 0.0000 |
| m16 | 2 | 13 | 22.5604 | 0.0002 | 50.6767 | 17.5323 | 0.0002 | 0.0000 |
| m55 | 3 | 12 | 25.7012 | 0.0003 | 51.5046 | 18.3603 | 0.0001 | 0.0000 |
| m73 | 3 | 12 | 25.7072 | 0.0003 | 51.5107 | 18.3664 | 0.0001 | 0.0000 |
| m32 | 3 | 12 | 25.7991 | 0.0002 | 51.6026 | 18.4582 | 0.0001 | 0.0000 |
| m93 | 4 | 11 | 28.6659 | 0.0004 | 52.1831 | 19.0388 | 0.0001 | 0.0000 |
| m56 | 2 | 13 | 24.4300 | 0.0001 | 52.5463 | 19.4020 | 0.0001 | 0.0000 |
| m17 | 2 | 13 | 24.7225 | 0.0001 | 52.8388 | 19.6945 | 0.0001 | 0.0000 |
| m94 | 3 | 12 | 27.3947 | 0.0001 | 53.1982 | 20.0539 | 0.0000 | 0.0000 |
| m42 | 4 | 11 | 29.7428 | 0.0002 | 53.2600 | 20.1157 | 0.0000 | 0.0000 |
| m80 | 5 | 10 | 32.3581 | 0.0003 | 53.6152 | 20.4709 | 0.0000 | 0.0000 |
| m74 | 3 | 12 | 27.8694 | 0.0001 | 53.6728 | 20.5285 | 0.0000 | 0.0000 |
| m43 | 4 | 11 | 30.3357 | 0.0002 | 53.8530 | 20.7086 | 0.0000 | 0.0000 |
| m18 | 1 | 14 | 23.4514 | 0.0000 | 53.9075 | 20.7632 | 0.0000 | 0.0000 |
| m81 | 5 | 10 | 32.9510 | 0.0003 | 54.2082 | 21.0638 | 0.0000 | 0.0000 |
| m4 | 3 | 12 | 28.7642 | 0.0001 | 54.5676 | 21.4233 | 0.0000 | 0.0000 |
| m75 | 2 | 13 | 26.5982 | 0.0000 | 54.7145 | 21.5702 | 0.0000 | 0.0000 |
| m5 | 3 | 12 | 29.3571 | 0.0001 | 55.1606 | 22.0162 | 0.0000 | 0.0000 |
| m61 | 4 | 11 | 31.9110 | 0.0001 | 55.4282 | 22.2839 | 0.0000 | 0.0000 |
| m62 | 4 | 11 | 32.5039 | 0.0001 | 56.0212 | 22.8769 | 0.0000 | 0.0000 |
| m44 | 3 | 12 | 30.6338 | 0.0000 | 56.4373 | 23.2929 | 0.0000 | 0.0000 |
| m82 | 4 | 11 | 33.2491 | 0.0001 | 56.7663 | 23.6220 | 0.0000 | 0.0000 |
| m46 | 4 | 11 | 33.6379 | 0.0000 | 57.1551 | 24.0108 | 0.0000 | 0.0000 |
| m35 | 3 | 12 | 31.9086 | 0.0000 | 57.7120 | 24.5677 | 0.0000 | 0.0000 |
| m6 | 2 | 13 | 29.6552 | 0.0000 | 57.7715 | 24.6271 | 0.0000 | 0.0000 |
| m84 | 5 | 10 | 36.6026 | 0.0001 | 57.8598 | 24.7154 | 0.0000 | 0.0000 |
| m47 | 3 | 12 | 32.3668 | 0.0000 | 58.1702 | 25.0259 | 0.0000 | 0.0000 |
| m8 | 3 | 12 | 32.6593 | 0.0000 | 58.4628 | 25.3184 | 0.0000 | 0.0000 |
| m63 | 3 | 12 | 32.8020 | 0.0000 | 58.6055 | 25.4612 | 0.0000 | 0.0000 |
| m85 | 4 | 11 | 35.3315 | 0.0000 | 58.8487 | 25.7044 | 0.0000 | 0.0000 |
| m65 | 4 | 11 | 35.8061 | 0.0000 | 59.3234 | 26.1790 | 0.0000 | 0.0000 |
| m9 | 2 | 13 | 31.3881 | 0.0000 | 59.5044 | 26.3601 | 0.0000 | 0.0000 |
| m36 | 3 | 12 | 34.0707 | 0.0000 | 59.8742 | 26.7299 | 0.0000 | 0.0000 |
| m66 | 3 | 12 | 34.5350 | 0.0000 | 60.3384 | 27.1941 | 0.0000 | 0.0000 |
| m57 | 3 | 12 | 34.6771 | 0.0000 | 60.4806 | 27.3363 | 0.0000 | 0.0000 |

| | | | | | | | | |
|---|---|---|---|---|---|---|---|---|
| m37 | 2 | 13 | 32.7996 | 0.0000 | 60.9159 | 27.7715 | 0.0000 | 0.0000 |
| m95 | 4 | 11 | 37.6418 | 0.0000 | 61.1591 | 28.0147 | 0.0000 | 0.0000 |
| m23 | 4 | 11 | 38.1124 | 0.0000 | 61.6296 | 28.4853 | 0.0000 | 0.0000 |
| m19 | 2 | 13 | 33.6985 | 0.0000 | 61.8148 | 28.6705 | 0.0000 | 0.0000 |
| m24 | 4 | 11 | 38.7053 | 0.0000 | 62.2225 | 29.0782 | 0.0000 | 0.0000 |
| m76 | 3 | 12 | 36.8453 | 0.0000 | 62.6488 | 29.5045 | 0.0000 | 0.0000 |
| m45 | 4 | 11 | 40.8809 | 0.0000 | 64.3982 | 31.2538 | 0.0000 | 0.0000 |
| m83 | 5 | 10 | 43.4962 | 0.0000 | 64.7534 | 31.6090 | 0.0000 | 0.0000 |
| m25 | 3 | 12 | 39.0034 | 0.0000 | 64.8068 | 31.6625 | 0.0000 | 0.0000 |
| m27 | 4 | 11 | 42.0075 | 0.0000 | 65.5247 | 32.3804 | 0.0000 | 0.0000 |
| m7 | 3 | 12 | 39.9023 | 0.0000 | 65.7058 | 32.5614 | 0.0000 | 0.0000 |
| m28 | 3 | 12 | 40.7363 | 0.0000 | 66.5398 | 33.3955 | 0.0000 | 0.0000 |
| m64 | 4 | 11 | 43.0491 | 0.0000 | 66.5664 | 33.4221 | 0.0000 | 0.0000 |
| m38 | 3 | 12 | 43.0467 | 0.0000 | 68.8502 | 35.7058 | 0.0000 | 0.0000 |
| m26 | 4 | 11 | 49.2505 | 0.0000 | 72.7677 | 39.6234 | 0.0000 | 0.0000 |
| m48 | 4 | 11 | 67.7711 | 0.0000 | 91.2884 | 58.1441 | 0.0000 | 0.0000 |
| m86 | 5 | 10 | 70.7358 | 0.0000 | 91.9930 | 58.8487 | 0.0000 | 0.0000 |
| m10 | 3 | 12 | 66.7925 | 0.0000 | 92.5960 | 59.4517 | 0.0000 | 0.0000 |
| m39 | 6 | 9 | 73.6769 | 0.0000 | 92.6996 | 59.5553 | 0.0000 | 0.0000 |
| m77 | 7 | 8 | 76.2922 | 0.0000 | 93.1057 | 59.9614 | 0.0000 | 0.0000 |
| m67 | 4 | 11 | 69.9394 | 0.0000 | 93.4566 | 60.3123 | 0.0000 | 0.0000 |
| m1 | 5 | 10 | 72.6982 | 0.0000 | 93.9554 | 60.8111 | 0.0000 | 0.0000 |
| m58 | 6 | 9 | 75.8451 | 0.0000 | 94.8678 | 61.7235 | 0.0000 | 0.0000 |
| m29 | 4 | 11 | 76.1407 | 0.0000 | 99.6580 | 66.5136 | 0.0000 | 0.0000 |
| m20 | 6 | 9 | 82.0464 | 0.0000 | 101.0692 | 67.9248 | 0.0000 | 0.0000 |

Model codes correspond to diagrams presented in Fig. S2. For each model we report the k, q, C-statistic (C), p-value, CICc value, ΔCICc value, l, and CICc weights (ω). P-values of the C-statistic, where significant indicate the model is rejected by the data. Models were based on 186 avian species.
